# A Novel L-Serine Exporter and Its Positive Regulator in *Corynebacterium glutamicum*

**DOI:** 10.1101/639088

**Authors:** Xiaomei Zhang, Yujie Gao, Ziwei Chen, Guoqiang Xu, Xiaojuan Zhang, Hui Li, Jinsong Shi, Mattheos Koffas, Zhenghong Xu

## Abstract

Exporters play an essential role in the fermentative production of amino acids. In *Corynebacterium glutamicum*, ThrE, which can export L-threonine and L-serine, is the only identified L-serine exporter so far. In this study, a novel L-serine exporter NCgl0580 was identified and characterized in *C. glutamicum* ΔSSAAI (SSAAI), and named as SerE (encoded by *serE*). Deletion of *serE* in SSAAI led to a 56.5% decrease in L-serine titer, whereas overexpression of *serE* compensated for the lack of *serE* with respect to L-serine titer. A fusion protein with SerE and enhanced green fluorescent protein (EGFP) was constructed to confirm that SerE localized at the plasma membrane. The function of SerE was studied by peptide feeding approaches, and the results showed that SerE is a novel exporter for L-serine and L-threonine in *C. glutamicum*. Subsequently, the interaction of a known L-serine exporter ThrE and SerE was studied, and the results suggested that SerE is more important than ThrE in L-serine export in SSAAI. Probe plasmid and electrophoretic mobility shift assays (EMSA) revealed NCgl0581 as the transcriptional regulator of SerE. Comparative transcriptomics between SSAAI and the NCgl0581 deletion strain showed that NCgl0581 regulated the transcription of 115 genes in *C. glutamicum*, among which the transcriptional level of NCgl0580 decreased 280-fold in a NCgl0581 deletion strain, indicating that NCgl0581 is a positive regulator of SerE. Thus, this study provides a novel target for L-serine and L-threonine export engineering as well as a novel global transcriptional regulator NCgl0581 in *C. glutamicum*.

**Importance:** Exporters are gaining increasing attention for improving industrial production of amino acids. This study identified a novel exporter NCgl0580 for L-serine and L-threonine in *C. glutamicum*, and its positive regulator (NCgl0581), which was shown to be a novel global transcriptional regulator in *C. glutamicum*. This study provides a new target for engineering efflux of L-serine and L-threonine, expands the exporter and transcriptional regulator family, and enriches our understanding of amino acid transport system in *C. glutamicum.*

## INTRODUCTION

L-serine has been identified as one of the top 30 most interesting building blocks for a range of chemicals and materials, and is used in cosmetic, pharmaceutical, and food industries (1, 2). Metabolic engineering of *Corynebacterium glutamicum* (*C.glutamicum*) for L-serine production has been focused on its terminal synthesis pathways and degradation pathways, and proven to be very useful for improving L-serine production in this organism (3-6); however, the L-serine productivity is still low for large-scale L-serine production.

Efflux is often overlooked as a bottleneck in metabolic pathways, and amino acid efflux transporters are increasingly gaining attention for optimizing industrial production of amino acids (7, 8). In recent decades, a number of export systems have been identified for excreting amino acids, such as L-lysine, L-cysteine, L-glutamate, L-threonine, L-arginine, L-methionine, and branched-chain amino acids, in *C. glutamicum* and *Escherichia coli* (8-14). However, to the best of our knowledge, except for ThrE (L-threonine and L-serine exporter) (12, 15), no other L-serine exporters have been reported in *C. glutamicum* so far. In *E. coli*, Mundhada et al. found that intracellular L-serine accumulation was toxic to the engineered strain modified to produce L-serine, and that following overexpression of *eamA* (which encodes L-cysteine exporter in *E. coli*), the engineered strain exhibited increased tolerance toward L-serine with higher L-serine productivity (1). Therefore, L-serine exporter in *C. glutamicum* could be a potential target for strain optimization to further improve L-serine production.

It has been reported that homologs similar to the exporters in *E. coli* might fulfil a comparable function in *C. glutamicum* (14, 16, 17). Accordingly, we hypothesized that the homolog to *eamA* (L-serine exporter in *E. coli*) (1) might be involved in L-serine export in *C. glutamicum*. In the present study, three homologs to *eamA*, namely, NCgl2050, NCgl2065, and NCgl0580, were determined, and their functions were identified by targeted gene deletion, respectively. The results showed that one of the genes, NCgl0580, was involved in L-serine export. Subsequently, localization and function of NCgl0580 were investigated, and the interaction of a known L-serine exporter ThrE (encoded by *thrE*) and the novel exporter NCgl0580 was studied. Furthermore, the transcriptional regulator of NCgl0580 was identified and studied.

## RESULTS

### Exploring putative L-serine exporters in *C. glutamicum*

In past studies, homologs of *E. coli* exporters have been shown to have similar functions in *C. glutamicum* (14, 16, 17). Therefore, we hypothesized that the *C. glutamicum* homolog to *eamA* (L-serine exporter in *E. coli*) (1) might be involved in L-serine export in this organism. According to the NCBI database, EamA belongs to the RhaT superfamily, and 15 records of related proteins associated with RhaT superfamily in *C. glutamicum* ATCC13032 were obtained. After eliminating duplicate records, three related genes, NCgl2050, NCgl2065, and NCgl0580, were obtained, which might be involved in L-serine export in *C. glutamicum*.

To verify the function of these putative proteins in *C. glutamicum* SSAAI (SSAAI), NCgl2050, NCgl2065, and NCgl0580 were respectively deleted in this strain. The results showed that the deletion of NCgl2050 and NCgl2065 did not produce any changes in cell growth and L-serine titer (Fig. 1A and 1B). Strikingly, deletion of NCgl0580 significantly reduced the L-serine titer in SSAAI, but did not affect the growth of the strain (Fig. 1C). SSAAI ΔNCgl0580 produced 11.31 g/L L-serine, which was 56.5% lower than that noted in SSAAI (Fig. 1C). However, plasmid-borne overexpression of NCgl0580 compensated for the lack of NCgl0580 with respect to L-serine titer, resulting in 26.76 g/L L-serine titer, similar to that generated by the parent strain SSAAI (Fig. 1D). As shown in Fig. 1D, when compared with SSAAI, the strain harboring the plasmid grew slowly to some extent in the logarithmic growth phase, finally reaching similar cell growth to SSAAI. This finding suggested that NCgl0580 might act as the L-serine exporter in *C. glutamicum*, and was named as SerE and its function was further investigated.

**FIG 1.**
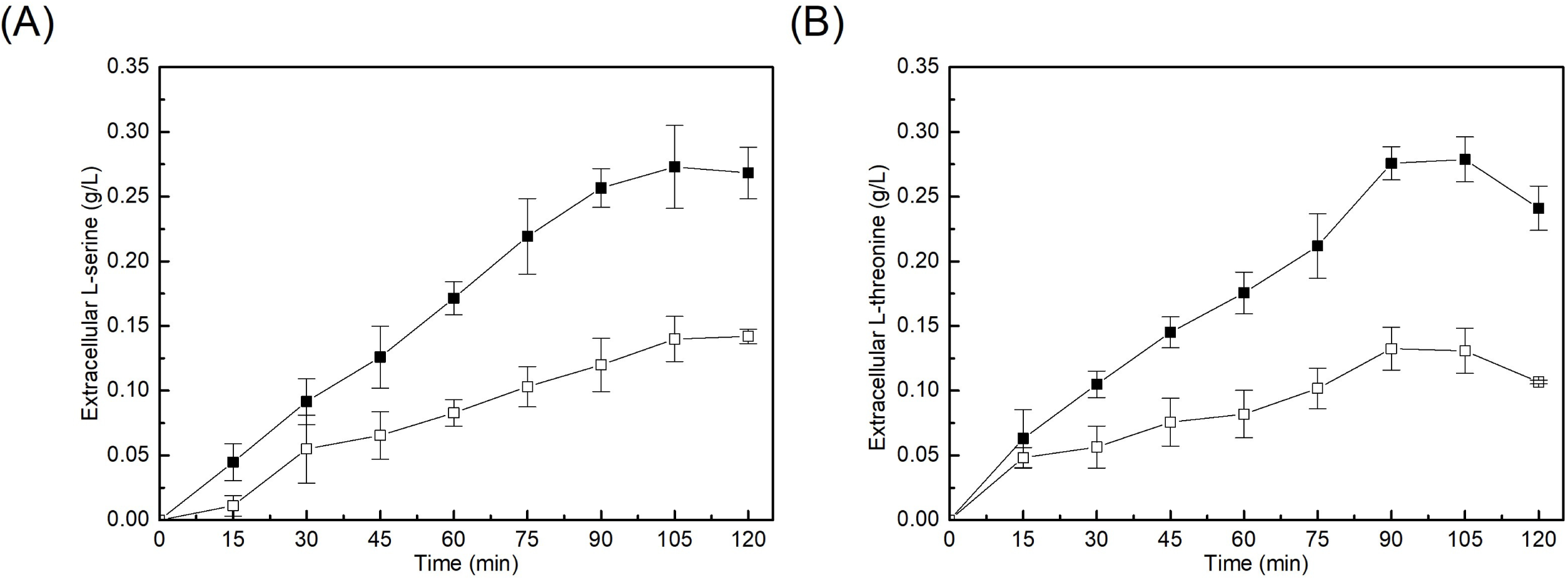
Effect of NCgl2050, NCgl2065, NCgl0580 deletion and plasmid compensating strain on SSAAI. **(A)** SSAAIΔNCgl2050 (Open symbols), SSAAI (Solid symbols). **(B)** SSAAIΔNCgl2065 (Open symbols), SSAAI (Solid symbols). **(C)** SSAAIΔNCgl0580 (Open symbols), SSAAI (Solid symbols). **(D)** The plasmid compensating strain SSAAIΔN0580-NCgl0580 (Open symbols). SSAAI (Solid symbols). Squares and circles indicate growth OD_562_ and L-serine titer, respectively. Three parallel experiments were performed. Error bars indicate standard deviations of results from three parallel experiments.

### Localization and function of SerE

According to the NCBI, SerE was presumed to be a hypothetical membrane protein of 301 amino acids, similar to permease of the drug/metabolite transporter (DMT) superfamily. The transmembrane helices of SerE were predicted by TMHMM Server v. 2.0, and SerE exhibited ten transmembrane-spanning helices with both amino- and carboxy-terminal ends in the cytoplasm.

To confirm the localization of SerE, SerE-EGFP fusion protein was expressed in SSAAI. Confocal microscopic observations of SSAAI-*egfp* and SSAAI-*serE*-*egfp* confirmed that EGFP and SerE-EGFP fusion proteins were successfully expressed, respectively (Fig.S1). To further verify the localization of SerE, membrane and cytoplasmic proteins from these two strains were extracted by ultrasonication, and the fluorescence of these proteins was determined using a fluorescence spectrophotometer. The fluorescence of the cytoplasmic proteins of SSAAI-*egfp* and membrane proteins of SSAAI-*serE*-*egfp* (Fig. 2) affirmed that SerE was localized at the plasma membrane in SSAAI.

**FIG 2.**
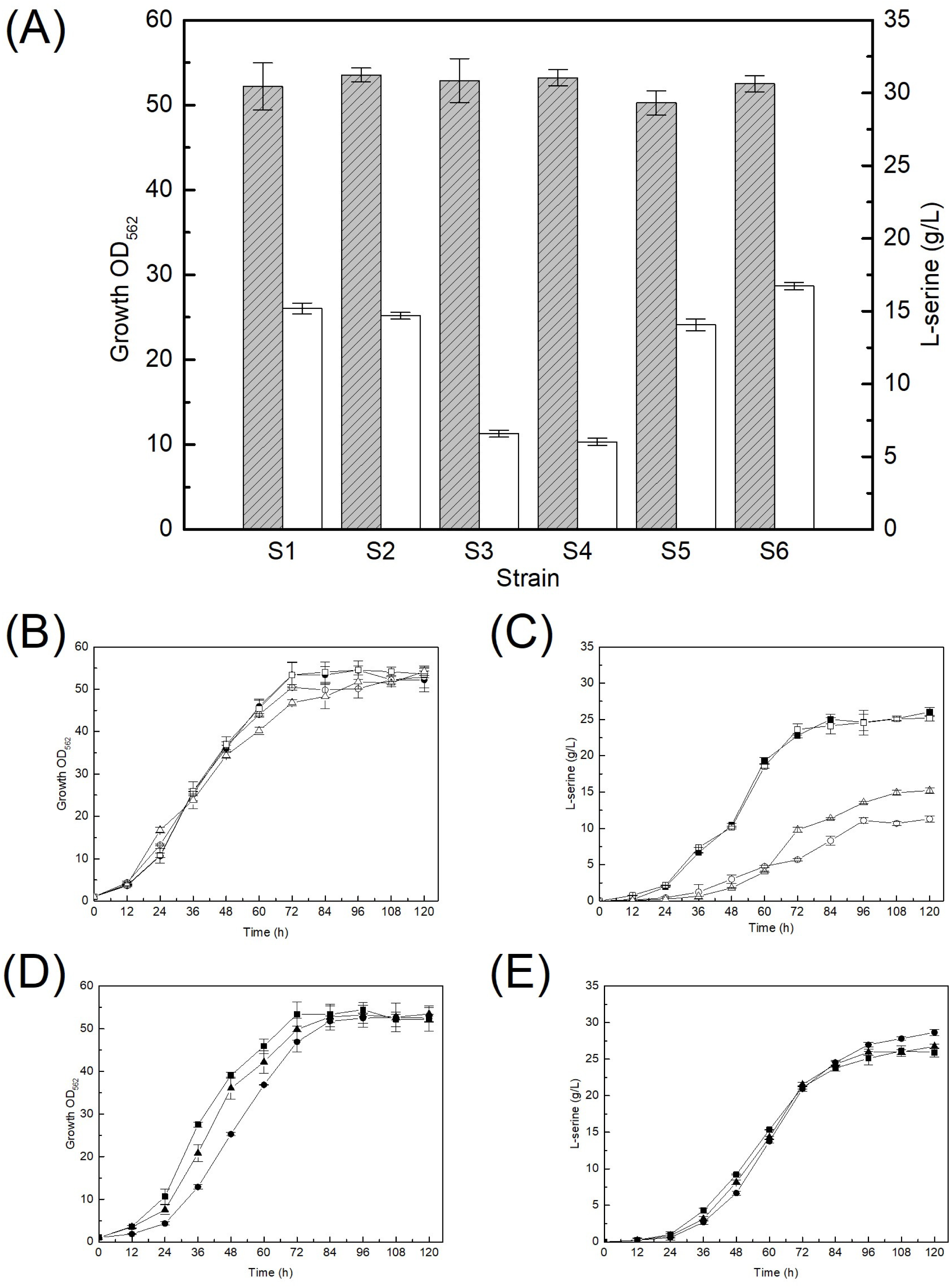
The fluorescence of cytoplasmic proteins and membrane proteins. The fluorescence of cytoplasmic proteins and membrane proteins of SSAAI-10 (SSAAI harboring plasmid pDXW-10 only, grey bar with slash), SSAAI-*egfp* (SSAAI expressing EGFP protein with pDXW-10, Grey bar) and SSAAI-*serE*-*egfp* (SSAAI expressing SerE-EGFP fusion protein with pDXW-10, White bar). Three parallel experiments were performed. Error bars indicate standard deviations of results from three parallel experiments.

To substantiate the function of SerE, a peptide feeding approach was employed by incubating SSAAI and SerE deletion strain, SSAAI Δ*serE,* with 2 mM of the dipeptide Ser-Ser, respectively, and measuring the concentration of extracellular L-serine. As shown in Fig. 3A, a higher L-serine concentration was detected in SSAAI, when compared with that in SSAAI Δ*serE*, confirming that SerE is a novel exporter of L-serine in *C. glutamicum*.

**FIG 3.**
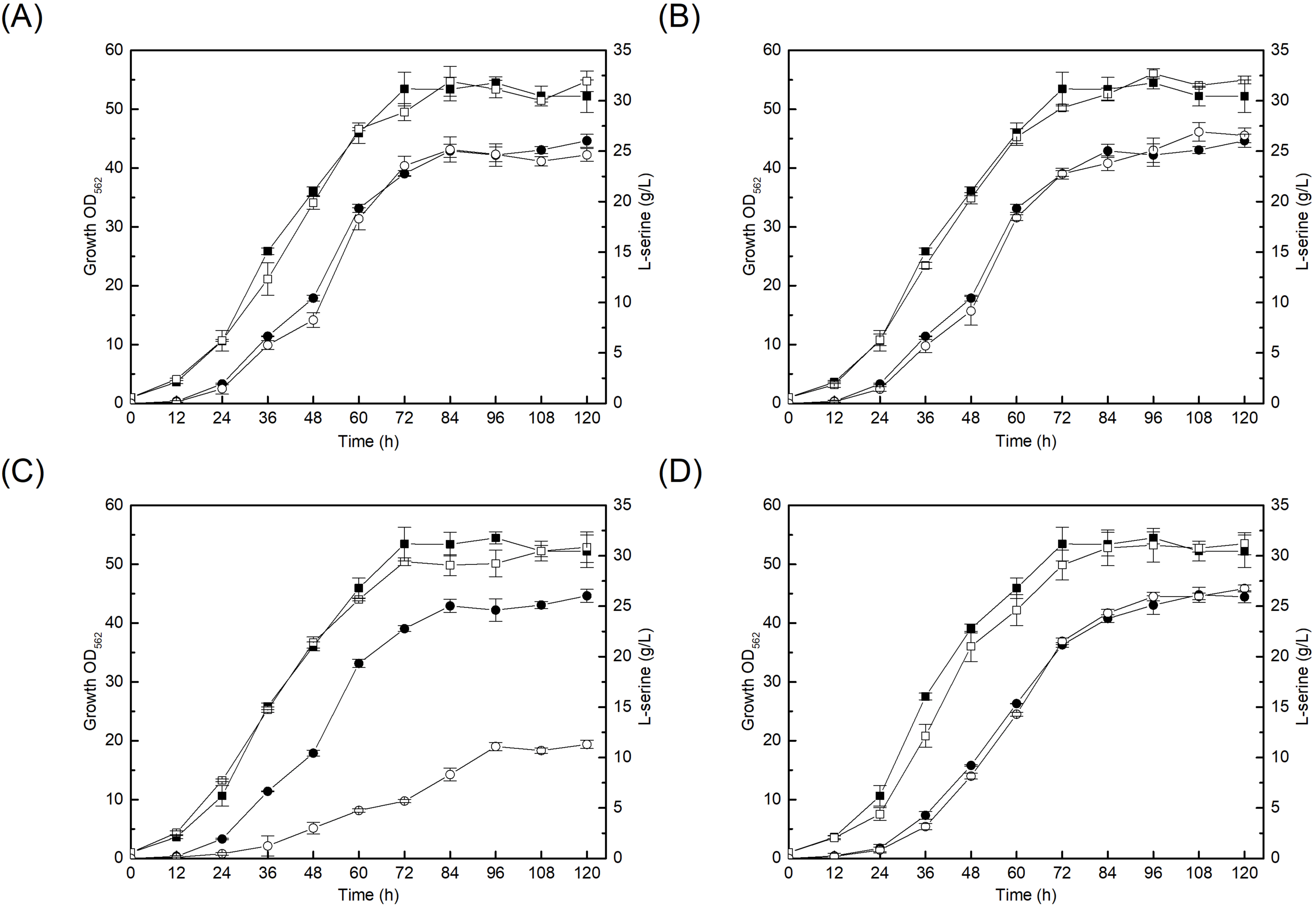
The result of amino acid export of SerE by using peptide feeding approach in SSAAI. **(A)** Extracellular concentration of L-serine in SSAAI (Solid symbols) and *serE* deletion strain SSAAI Δ*serE* (Open symbols). **(B)** Extracellular concentration of L-threonine in SSAAI (Squares) and *serE* deletion strain SSAAI Δ*serE* (Circles). Three parallel experiments were performed. Error bars indicate standard deviations of results from three parallel experiments.

It is known that L-cysteine uptake system in *E. coli* (encoded by *eamA*) also catalyzes L-serine uptake (1), and that L-threonine exporter in *C. glutamicum* (encoded by *thrE*) also transports L-serine (12). We therefore analyzed whether the novel exporter SerE could uptake L-cysteine and L-threonine. The export experiments with dipeptides (Thr-Thr, Cys-Cys) were performed using SSAAI and SSAAI Δ*serE*. The dipeptides were added at a concentration of 2 mM to the medium, and the extracellular amino acid concentrations at different time intervals were determined by HPLC. The results revealed that the concentration of L-cysteine was comparable in both strains and did not significantly change (data not shown), indicating that SerE does not export L-cysteine. Interestingly, the concentrations of L-threonine in SSAAI Δ*serE* were lower than those in SSAAI (Fig. 3B), indicating that SerE is also an exporter of L-threonine in *C. glutamicum*.

### Interaction of ThrE and SerE

It is well known that *thrE* encodes ThrE that can export L-threonine and L-serine in *C. glutamicum* ATCC13032 (12). To understand the interaction of ThrE and SerE on L-serine uptake, *thrE* was deleted in SSAAI (SSAAI Δ*thrE),* and no significantly change was observed in L-serine uptake for the deletion mutant (Fig. 4A). In contrast, deletion of SerE significantly reduced the L-serine titer in SSAAI, and resulted in a slight change in cell growth (Fig. 1C). The SSAAI Δ*serE* produced 11.31 g/L L-serine, which was 56.5% lower than that produced by SSAAI (Fig. 1C). Subsequently, *thrE* and *serE* double deletion mutant was constructed, which exhibited cell growth comparable to that of SSAAI (Fig. 4B), and produced 10.34 g/L L-serine, which was 60% lower than that observed in SSAAI (Fig. 4C).

**FIG 4.**
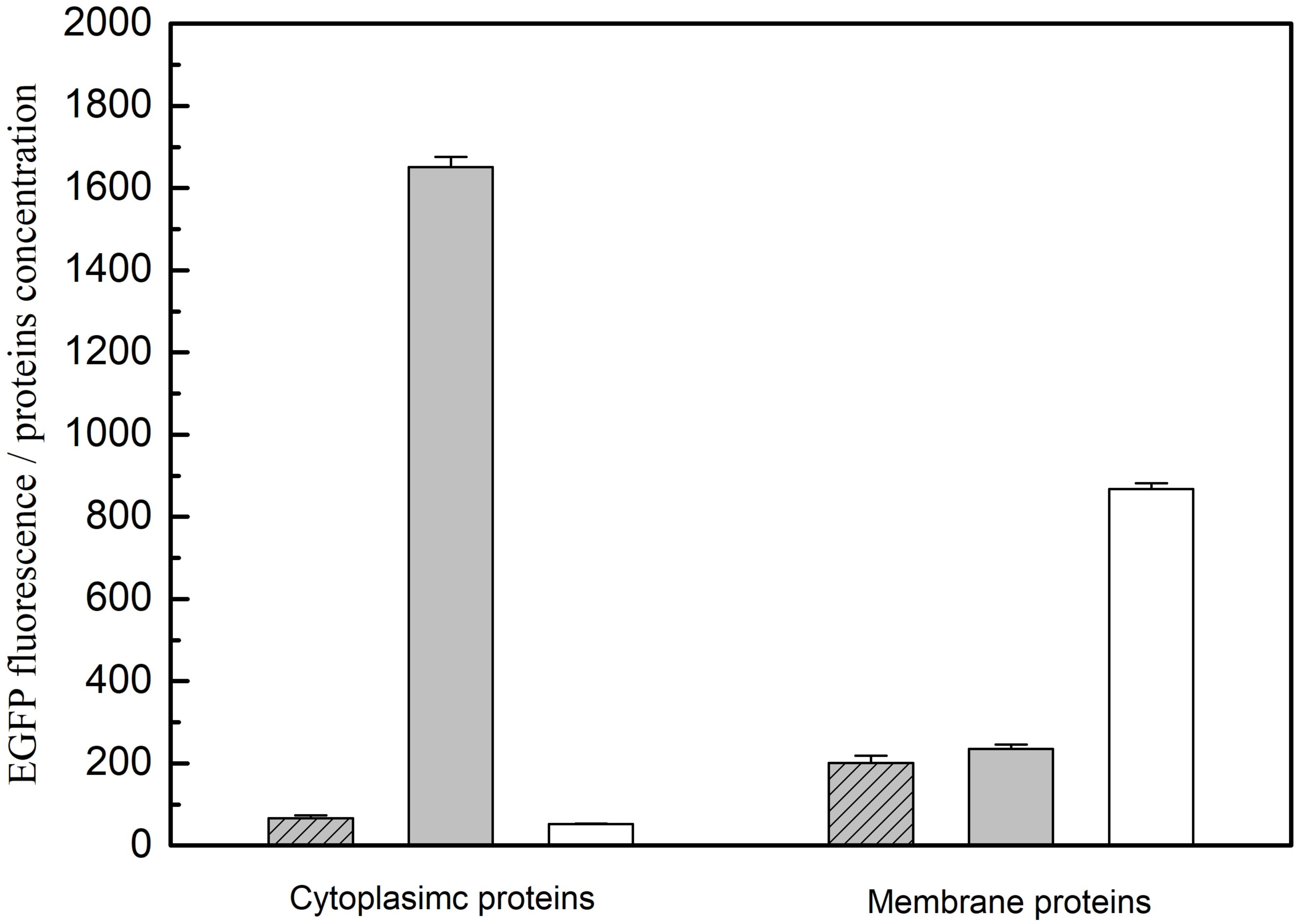
The effect of *thrE, serE* deletion or overexpression on SSAAI. **(A)** The cell growth (Grey bar with slash), L-serine titer (White bar). SSAAI (S1), *thrE* deletion strain (S2), *serE* deletion strain (S3), *thrE* and *serE* deletion strain (S4), *thrE* overexpression strain (S5) and *serE* overexpression strain (S6). **(B)** Growth OD_562_ of stain with *thrE* and *serE* double deletion, SSAAI (Solid squares), SSAAI Δ*thrE*-Δ*serE* (Open triangles), SSAAI Δ*thrE* (Open squares) and SSAAI *ΔserE* (Open circles). **(C)** L-serine titer of stain with *thrE* and *serE* double deletion, SSAAI (Solid circles), SSAAI Δ*thrE*-Δ*serE* (Open triangles), SSAAI Δ*thrE* (Open squares) and SSAAI Δ*serE* (Open circles). **(D)** Growth OD_562_ of strains with *serE* overexpression, SSAAI-10 (Squares), SSAAI-*serE* (Circles), and SSAAI Δ*serE*-*serE* (Triangles). **(E)** L-serine titer of strains with *serE* overexpression, SSAAI-10 (Squares), SSAAI-*serE* (Circles), and SSAAIΔ*serE*-*serE* (Triangles). Three parallel experiments were performed. Error bars indicate standard deviations of results from three parallel experiments.

Furthermore, *thrE* and *serE* were overexpressed in SSAAI to obtain SSAAI-*thrE* and SSAAI-*serE* respectively. While L-serine accumulation in SSAAI-*thrE* was similar to that in SSAAI (Fig. 4A), the production of L-serine in SSAAI-*serE* reached 28.67 g/L, which was 10.5% higher than that noted in SSAAI (Fig.4D). However, a decrease in cell growth was observed in SSAAI-*serE* before 72 h of fermentation, when compared with that found in SSAAI (Fig.4E). These results suggested that SerE is more important than ThrE in L-serine export in SSAAI.

### Transcriptional regulator of SerE

The gene NCgl0581, located upstream of *serE* and divergently transcribed from *serE* (Fig. S2), and its product (consisting of 303 amino acids) were found to be members of the LysR-type transcriptional regulators (LTTRs) family. It must be noted that LTTRs were initially described as regulators of divergently transcribed genes (18). In a previous study on *C. glutamicum,* LysG, located upstream of L-lysine exporter gene *lysE,* was observed to encode a LysR-type transcriptional regulator, confirming that LysG is a positive transcriptional regulator of *lysE* (19). Accordingly, we speculated that NCgl0581 might be involved in the control of *serE* transcription.

To determine the function of NCgl0581, a mutant strain with NCgl0581 deletion was constructed. As shown in Fig. 5, the growth of SSAAI ΔNCgl0581 was similar to that of the parent strain SSAAI. However, the L-serine titer of SSAAI ΔNCgl0581 was 11.08 g/L, which was 57.4% lower than that of the parent strain, indicating that NCgl0581 played an important role in L-serine production. Subsequently, the effect of NCgl0581 on *serE* expression was further investigated by using the probe plasmid pDXW-11. Two recombinant strains, SSAAI ΔNCgl0581-1 (harboring the plasmid pDXW-11-1, Fig. S3A) and SSAAI ΔNCgl0581-0 (harboring the plasmid pDXW-11-0, Fig. S3B) were constructed, and their fluorescence during fermentation was measured. The fluorescence of SSAAI ΔNCgl0581-1 was stronger than that of SSAAI ΔNCgl0581-0 during the fermentation process (Fig. S3C), revealing that NCgl0581 functioned as a positive regulator of *serE* expression. To verify whether the regulatory protein NCgl0581 binds to the upstream region of SerE, electrophoretic mobility shift assay (EMSA) was performed by using the DNA probe labeled with biotin, and the result clearly indicated that NCgl0581 binds to this region (Fig. S4).

**FIG 5.**
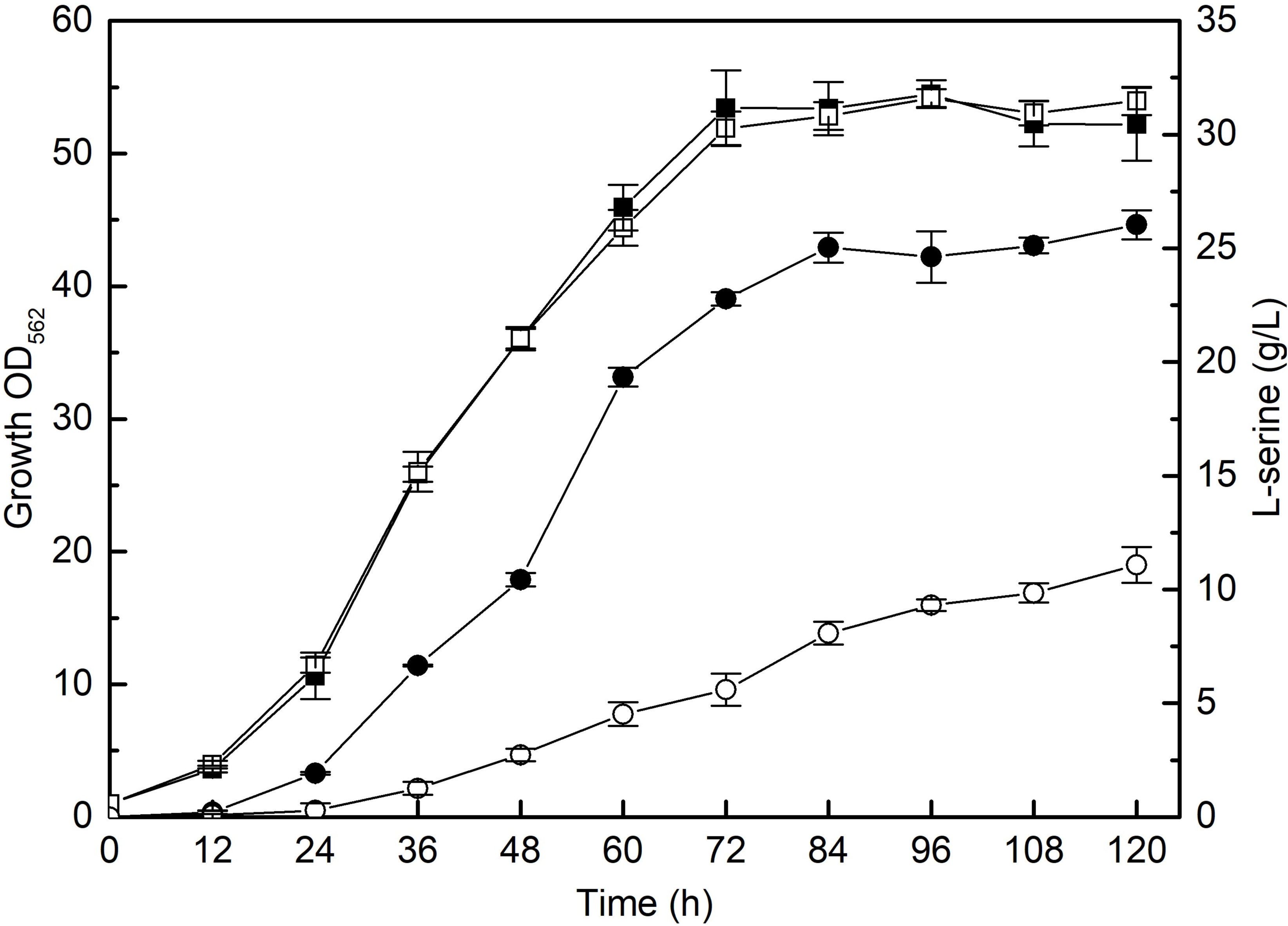
Effect of NCgl0581 deletion on SSAAI. The cell growth (Squares) and L-serine titer (Circles) of SSAAI (Solid symbols) and SSAAIΔNCgl0581 (Open symbols), respectively. Three parallel experiments were performed. Error bars indicate standard deviations of results from three parallel experiments.

To confirm whether NCgl0581 is a specific regulator of SerE, transcriptome sequencing was performed using SSAAI and NCgl0581 deletion strain. The findings showed that the transcription levels of 115 genes were altered, including 56 genes upregulated and 59 genes downregulated, in response to NCgl0581 deletion, indicating that NCgl0581 is a global transcriptional regulator in *C. glutamicum.* The genes with significant transcriptional change (≥4-fold) are shown in Tables 2 and 3. The transcriptional level of *serE* was significantly decreased by 280-fold following NCgl0581 deletion, revealing that NCgl0581 is a positive regulator of *serE*. Furthermore, NCgl0581 deletion downregulated the two ABC transporter permeases (NCgl0638 and NCgl0484) and ABC transporter periplasmic component (NCgl0639) by 12-, 6.3-, and 7.5-fold, respectively, and upregulated ABC transporter periplasmic component (NCgl1405) by 5.88-fold, suggesting that NCgl0581 is involved in the synthesis of substances transported through ABC transporter.

**TABLE 1.**
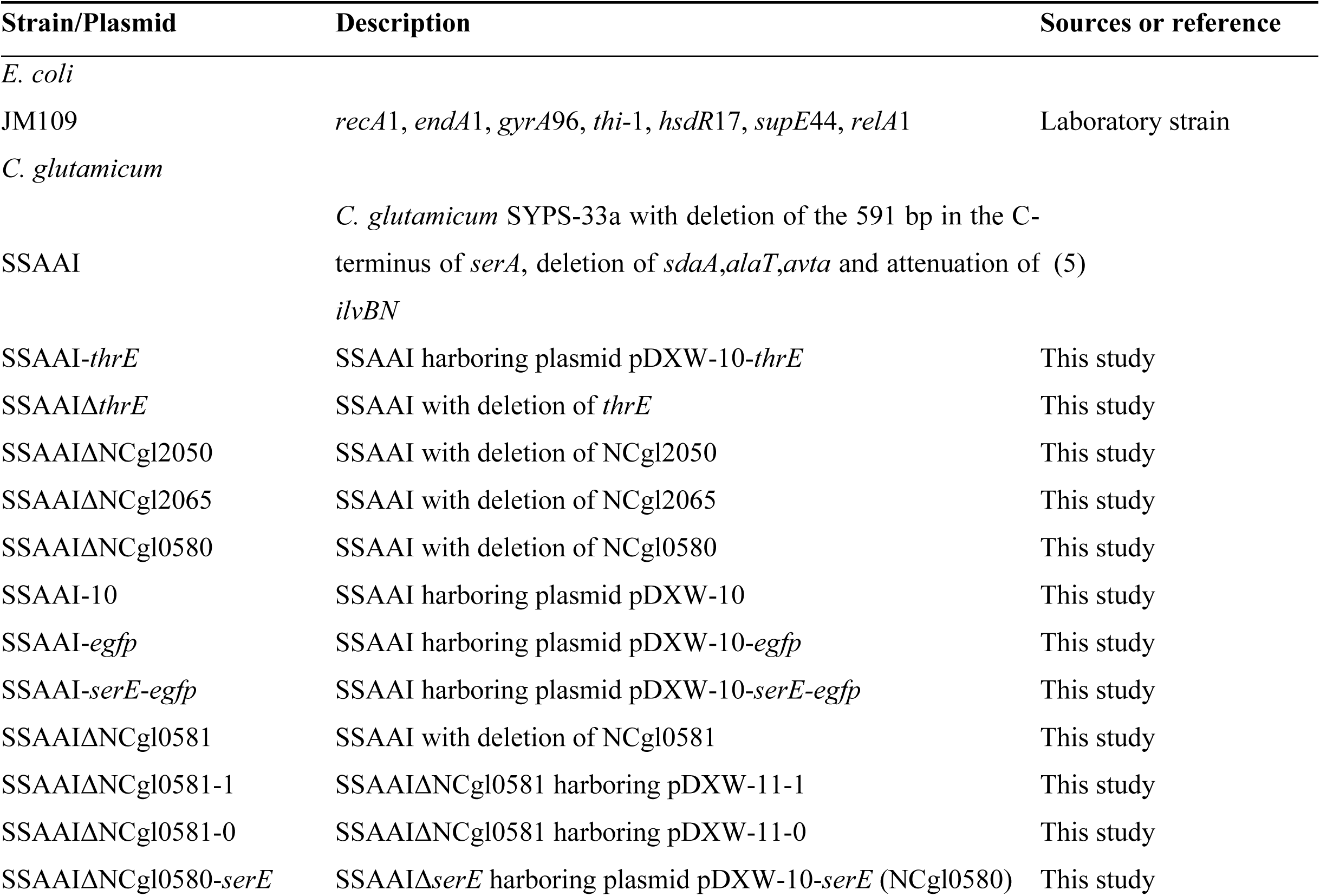
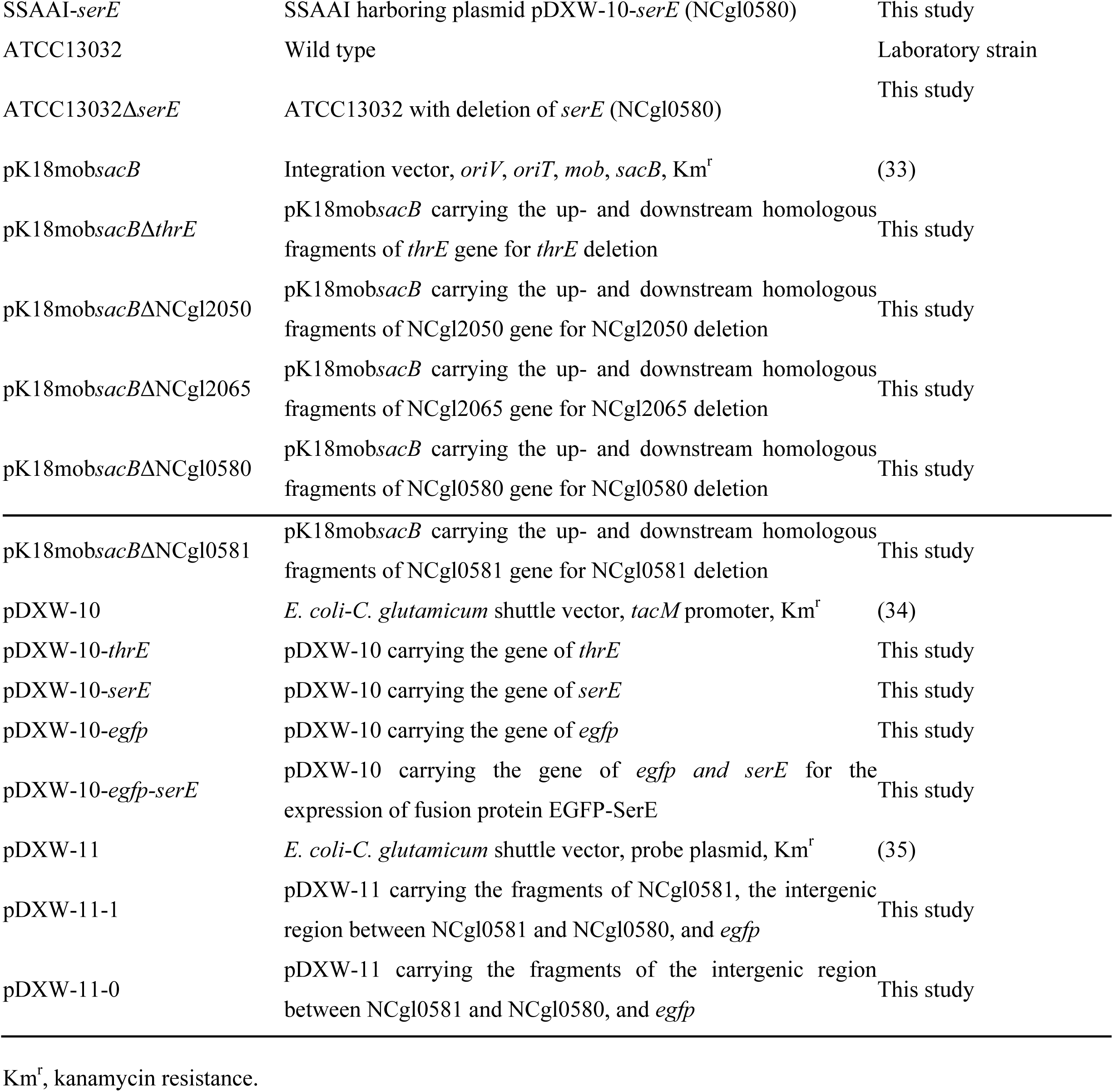
Strains and plasmids used in this study.

**TABLE 2.**
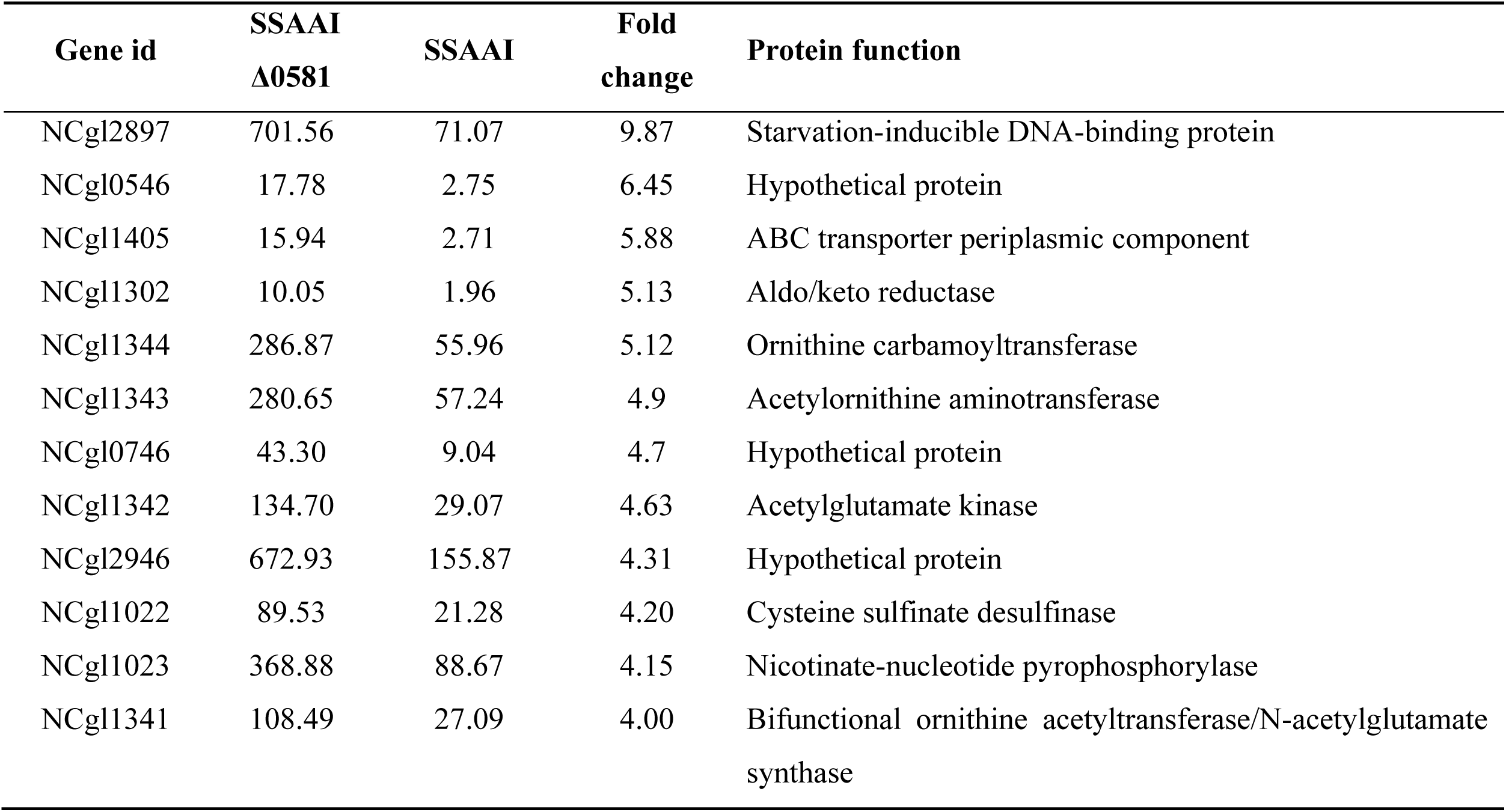
The genes significantly up regulated by NCgl0581 deletion.

**TABLE 3.** The genes significantly down regulated by NCgl0581 deletion.

| Gene id | SSAAI<br>$\Delta 0581$ | SSAAI | Fold<br>change | Protein function |
| --- | --- | --- | --- | --- |
| NCgl0580 | 18.40 | 5152.54 | 280.02 | Hypothetical protein |
| NCgl0638 | 1.71 | 20.97 | 12.22 | ABC transporter permease |
| NCgl0639 | 11.00 | 82.47 | 7.49 | ABC transporter periplasmic component |
| NCgl2943 | 207.03 | 1355.55 | 6.54 | Hypothetical protein |
| NCgl0943 | 16.19 | 103.52 | 6.39 | AraC family transcriptional regulator |
| NCgl0484 | 2.32 | 14.57 | 6.28 | ABC transporter permease |
| NCgl2942 | 283.52 | 1776.15 | 6.26 | NADH: flavin oxidoreductase |
| NCgl0166 | 13.41 | 79.70 | 5.94 | Hypothetical protein |
| NCgl0324 | 2.11 | 11.87 | 5.61 | Zn-dependent alcohol dehydrogenase |
| NCgl0282 | 5.19 | 28.25 | 5.44 | 4-Hydroxyphenyl-beta-ketoacyl-CoA hydrolase |
| NCgl1975 | 102.94 | 503.75 | 4.89 | Hypothetical protein |
| NCgl2893 | 1.25 | 6.08 | 4.84 | Efflux system protein |
| NCgl0155 | 9.11 | 43.69 | 4.79 | 5-Dehydro-2-deoxygluconokinase |
| NCgl0014 | 10.02 | 47.76 | 4.76 | Hypothetical protein |
| NCgl2953 | 7.68 | 35.80 | 4.66 | Sugar permease |
| NCgl2744 | 12.26 | 55.19 | 4.50 | Hypothetical protein |
| NCgl2970 | 15.22 | 67.51 | 4.43 | ABC transporter periplasmic component |
| NCgl0608 | 23.06 | 100.35 | 4.35 | ABC transporter permease |
| NCgl0258 | 4.51 | 19.50 | 4.32 | Arsenite efflux pump ACR3 |
| NCgl0281 | 16.83 | 67.69 | 4.02 | Dehydrogenase |

## DISCUSSION

Transport engineering is becoming an attractive strategy for strain improvement (8, 13, 14). However, only a relatively limited number of exporters of amino acids have been identified in *C. glutamicum* (Table S1) (9, 11-14, 20-24). In the present study, SerE was identified as a novel L-serine exporter in *C. glutamicum*. Further analysis showed that SerE could also uptake L-threonine (Fig. 3B), but not L-cysteine, similar to ThrE, which can export both L-serine and L-threonine in *C. glutamicum* (12). It is assumed that the presence of -OH in both L-serine and L-threonine might be the reason for these exporters to transport the two substrates. Based on homology search, SerE was found to be similar to a member of the DMT superfamily. Although DMT superfamily proteins are involved in the transport of a wide range of substrates, there are only a few reports available on their structures and mechanisms of substrate transport. Tsuchiya et al. performed structural and functional analyses of YddG, a DMT protein, and provided insight into the common transport mechanism shared among the DMT superfamily members (25). It has been reported that analyses of the crystal structure data of exporters could help to elucidate the elusive transport mechanism (26), and in the future, we plan to investigate further the SerE structures and mechanisms of substrate transport.

To explore the interaction between the known L-serine exporter ThrE and the novel exporter SerE on L-serine uptake, ThrE and SerE single and double mutants were constructed. The results showed that *serE* and *thrE* double deletion mutant could still accumulate 10.34 g/L L-serine (Fig. 4), suggesting that *C. glutamicum* might also possess other L-serine exporter systems. The evolution of multiple exporter systems for a single substrate is beneficial for the survival of bacteria in variable environment (7, 27). It must be noted that overexpression of *serE* in SSAAI resulted in 10.5% increase in L-serine titer, but a decrease in cell growth. This could be due to the use of constitutive-type promoter to overexpress SerE, which resulted in higher L-serine efflux. As sufficient L-serine content is important to maintain cell growth, a decrease in cell growth was noted as a stress response to *serE* overexpression. In future studies, better tuning of the *serE* expression will be performed in SSAAI by testing different promoters and RBS.

NCgl0581 was identified as a SerE transcriptional regulator, and EMSA was performed to confirm the binding sites of NCgl0581 with the promoter of SerE. A previous study reported that the first member of the protein-gene pairs, ArgP-*argO* in *E. coli* and LysG-*lysE* in *C. glutamicum,* is an LysR-type transcriptional regulator, while the second member is its target gene encoding an amino acid exporter (19, 28, 29). Similarly, NCgl0581-*serE* might also be a protein-gene pair sharing the same regulation mechanism. A serine-biosensor based NCgl0581 was reported by Binder et al. (30), and based on this study, we constructed a biosensor for L-serine and found that NCgl0581 activated NCgl0580 (SerE) expression in the presence of L-serine, the expression of NCgl0580 enhancing with the L-serine titer increasing (31). However, NCgl0581 did not activate the expression of NCgl0580 in the presence of L-alanine and L-valine. To further confirm whether SerE could uptake L-alanine and L-valine, peptide feeding assays were employed using dipeptides (Ala-Ala, Val-Val) with SSAAI and SSAAI Δ*serE*. The results revealed that SerE could neither export L-alanine nor L-valine (data not shown). Moreover, transcriptome sequencing showed that NCgl0581 regulated 115 genes in *C. glutamicum*, suggesting that NCgl0581 was a novel global transcriptional regulator in *C. glutamicum*. Transcriptional regulators and their roles in expression control of target genes are important for metabolic engineering of *C. glutamicum* for industrial applications (32), and this study provided a new member of transcriptional regulator family.

In conclusion, a novel exporter SerE and its positive regulator NCgl0581 were identified in *C. glutamicum,* with SerE also exhibiting the ability to uptake L-threonine and NCgl0581 acting as a novel global transcriptional regulator in *C. glutamicum*. These results enrich our understanding of amino acid transport and can provide additional targets for exporter engineering in *C. glutamicum*.

## MATERIALS AND METHODS

### Strains, plasmids, and growth conditions

The strains and plasmids used in this study are listed in Table 1. *E. coli* JM109 was used as the cloning host, and was grown in lysogeny broth (LB) medium (containing 5.0 g/L yeast extract, 10.0 g/L tryptone, and 10.0 g/L NaCl) at 37°C and 220 rpm. The engineered SSAAI (CGMCC No.15170) was constructed in our laboratory by knocking out 591 bp of the C-terminal domain of *serA*, deleting *sdaA, avtA*, and *alaT*, as well as attenuating *ilvBN* in the genome of *C. glutamicum* SYPS-062-33a (CGMCC No. 8667). The seed and fermentation media for *C. glutamicum* were prepared as described previously (5). The *C. glutamicum* strains were pre-incubated in the seed medium overnight to an optical density (OD_562_) of about 25, and then inoculated at an initial concentration of OD_562_=1 into a 250 mL flask containing 25 mL of the fermentation medium at 30°C and 120 rpm. The antibiotic kanamycin (50 mg/L) was added when necessary. Samples were withdrawn periodically for the measurement of residual sugar, amino acids, and OD_562_ as described previously (5).

### Construction of plasmids and strains

The primers used in this study for gene expression/deletion are listed in Table S2. Gene deletion was performed using the nonreplicable deletion vector pK18mob*sacB*, as reported previously (33). For example, to achieve *thrE* deletion, the homologous-arm fragments for *thrE* deletion were amplified from SSAAI chromosome using the primer pairs *thrE*1/2 for the upstream fragment and *thrE*3/4 for the downstream fragment. Then, with the two fragments as templates, a crossover PCR was performed using the primer pair *thrE*1/4. The truncated product of *thrE* was digested with *Xba*I and *Hind*III and ligated to the vector pK18mob*sacB* that was similarly treated. The recombinant plasmid pK18mob*sacB*Δ*thrE* was transformed into SSAAI competent cells by electroporation, and chromosomal deletion was performed by selecting cells that were kanamycin resistant and sucrose nonresistant, and verified by PCR.

The pDXW-10 and pDXW-11 plasmids were used to overexpress genes in *C. glutamicum* (34, 35). The recombinant plasmids were constructed as follows: the genes *thrE* and *serE* were amplified, digested, and ligated to the pDXW-10 plasmid that was digested with *Hind*III/*Bgl*II. The plasmid harboring the fusion protein, SerE-EGFP (enhanced green fluorescent protein) was constructed by using the method reported in a previous study (16). To confirm the role of NCgl0581 on NCgl0580 expression, the fragment consisting of intergenic region of NCgl0581 and NCgl0580 and EGFP with or without NCgl0581 was ligated to the plasmid pDXW-11 by Clon Express MultiS One Step Cloning Kit (Vazyme, Nanjing, China). The strains were constructed by electroporation with the corresponding plasmids.

### Confocal microscopic observation

The strains SSAAI-10 (SSAAI harboring plasmid pDXW-10), SSAAI-*egfp*, and SSAAI-*serE*-*egfp* were grown in the seed medium and harvested during the exponential phase. The cells were washed twice and maintained in PBS (pH 7.4), mounted on a microscope slide, and observed under a Leica laser scanning confocal microscope (Leica, TCS SP8; Leica, Wetzlar, Germany) equipped with a HC PL Apo 63x/1.40 Oil CS2 oiL-Immersion objective, with excitation filter at 488 nm and emission filter at 510-550 nm. The digital images were acquired and analyzed with Lecia Application Suite X 2.0.

### Membrane and cytoplasmic protein extraction and fluorescence measurements

The strains SSAAI-10, SSAAI-*egfp*, and SSAAI-*serE*-*egfp* were used for extracting membrane and cytoplasmic proteins to determine SerE localization. The extraction was performed using Membrane and a Cytoplasmic Protein Extraction Kit according to the manufacturer’s protocol (Beyotime, Nanjing, China). The cells were washed twice with PBS (pH 7.4) and disrupted by ultrasonication on ice (pulse, 4 s; interval, 6 s; total duration, 30 min) (Sonics Vibra-Cell™, Sonics, Newtown, CT, USA). The supernatant containing cytoplasmic proteins was collected by centrifugation (700 × *g*, 4 °C for 10 min), and the precipitate was used for extracting membrane proteins. The protein concentration was determined with a Modified BCA Protein Assay Kit (Sangon, China). After extraction, the fluorescence intensity (excitation at 488 nm, emission at 517 nm) of the membrane and cytoplasmic proteins was determined using fluorescence spectrophotometer (Synergy H4; BioTek, Winooski, VT, USA).

### Amino acid export assay

For ascertaining the function of *serE*, a dipeptide Ser-Ser addition assay was conducted (12). In brief, pre-incubated cells (in seed medium) were washed once with CGXII minimal medium (36), inoculated into CGXII minimal medium with 2 mM Ser-Ser (other dipeptide), and incubated for 2 h at 30°C. Then, the cells were harvested, washed once with cold CGXII minimal medium, and resuspended in CGXII minimal medium. Amino acid excretion was initiated by adding 2 mM Ser-Ser (another dipeptide). HPLC was used to determine the concentrations of amino acids (16).

### Analytical procedures

Cell density (OD_562_) was measured using an AOE UV-1200S UV/vis spectrophotometer (AOE Instruments Co. Inc., Shanghai, China). Glucose concentration was determined using SBA-40E glucose analyzer (Biology Institute of Shandong Academy of Sciences, China). For measurement of extracellular L-serine concentration in shake-flask fermentation, 500 μL of the culture were centrifuged at 700 ×*g* for 5 min, and the supernatant was used for detection after appropriate dilution. To ascertain intracellular L-serine concentration, 300 μL of the culture were centrifuged at 700 ×*g* and 4°C for 10 min, and 300 μL of water were added to the cells. The cells were disrupted by FastPrep-24 5G instrument (5 m/s, 120 s, MP Biomedicals, Shanghai, China). The cytoplasmic volume was assumed to be 2 μL/mg dry cell weight (23). The titers of intracellular and extracellular L-serine and other amino acids were analyzed by HPLC using phenylisothiocyanate as a precolumn derivatization agent, according to our previously study (24).

### EMSA

To identify the binding site of NCgl0581 in the NCgl0580 promoter region, EMSA was conducted by using Non-Radioactive EMSA Kits with Biotin-Probes User’s Manual VER. 5.11 (Viagene Biotech Inc, Changzhou, China), according to the manufacturer’s instruction. The consensus oligonucleotides were BIO-JNZXM-TP (5’-AAACAGCCAA CTATAGTTAAGTAATA-3’) and BIO-JNZXM-BM (5’-TATTACTTAACTATAGTTGGCTGTTT-3’).

## ACKNOWLEDGMENTS

This work was financially supported by the National Natural Science Foundation of China (No. 31600044). Program of National First-class Discipline Program of Light Industry Technology and Engineering (LITE2018-11). Program of Introducing Talents of Discipline to Universities (No.111-2-06). International Joint Research Laboratory for Engineering Synthetic Biosystem for Intelligent Biomanufacturing at Jiangnan University. Top-notch Academic Programs Project of Jiangsu Higher Education Institutions (TAPP). And China Postdoctoral Science Foundation (No.2018M640452).

